# Evolution of symbiont transmission in conditional mutualisms in spatially and temporally variable environments

**DOI:** 10.1101/2020.05.05.079103

**Authors:** Alexandra Brown, Erol Akçay

## Abstract

Symbiotic relationships affect the fitness and organismal function of virtually all organisms. In many cases, the fitness effects of symbiosis may be beneficial or harmful depending on the environment. The hosts of such symbionts are favored to acquire them only when the symbiont is beneficial. However, it is not clear whether such selection favors vertical or horizontal transmission, both, or neither. To address this question, we model the evolution of transmission mode in a conditional mutualism experiencing spatial and temporal environmental variation. We find that when symbionts affect host lifespan, but not fecundity, horizontal transmission can contain them to beneficial environments. Vertical transmission can produce symbiont containment when the environmental state is synchronized across locations. We also find an emergent trade-off between horizontal and vertical transmission, suggesting that physiological constraints are not required for the evolution of limits on the total amount of transmission.

## 1 Introduction

Symbiosis is ubiquitous in our world. Virtually all multicellular organisms engage in it, and examples of both mutualistic and parasitic symbioses have been discovered in an enormous range of species. Many symboses are in fact neither purely parasitic nor purely mutualistic. The costs and benefits of these interactions instead depend on the environmental context (Bronstein, 1994; Hoeksema et al., 2010; Daskin and Alford, 2012; Chamberlain et al., 2014). For example, the bacterial symbiont *Hamiltonella defensa* protects its aphid host from parasitoid wasps and is highly beneficial when wasps are present (Oliver et al., 2003). However, in the absence of parasitoids, infection with *H. defensa* can have negative effects, including shortening aphid lifespan (Vorburger and Gouskov, 2011) and increasing susceptibility to predation by ladybugs (Polin et al., 2014).

Context-dependent symbioses have been found in a wide range of species, from symbioses between *Symbiodinium* dinoflagellates and corals (Baker et al., 2013) to those between intestinal nematodes and mice (Sutherland et al., 2011) and between trypanosomes and squirrels (Munger and Holmes, 1988). Bacteria are not just context-dependent symbionts but also hosts to their own conditionally mutualistic plasmids (Carroll and Wong, 2018), while fungi are engaged in a wide variety context-dependent interactions, including with grasses as fungal endophytes (Afkhami and Rudgers, 2009; Davitt et al., 2011; Yule et al., 2011; Cheplick et al., 1989), with plants as mycorrhizae (Johnson et al., 1997; Hoeksema et al., 2010; Heath and Tiffin, 2007), and with insect hosts, including bark beetles (Klepzig and Six, 2004) and black flies (McCreadie et al., 2005).

While extensive theory exists for the evolution of purely mutualistic and purely parasitic interactions, less is known about the evolution of context-dependent symbioses. With the many context-dependent interactions already known and likely waiting to be discovered, this is an important gap in our understanding of species interactions.

Previous theory on context-dependent symbioses focused on two main sets of questions. The first set considers the effects of context dependence in factors other than the cost and benefits of symbiosis. Theory in this set has investigated the effects of context-dependent transmission (Gundel et al., 2008), host birth rates (Ferris and Best, 2018), environmental productivity (Harrison et al., 2013; Poisot et al., 2012; Hochberg et al., 2000), and the specificity of selection (Mostowy and Engelstadter, 2011). The second category of models deals with context-dependence in the costs and benefits of symbiosis itself, more similar to the setting that we are considering. Vale et al. (2011) modeled a parasitic interaction where the costs of being parasitized depended on environmental conditions, and showed this affects the fraction of infected hosts and the opportunity for selection. Nuismer et al. (2003), Gomulkiewicz et al. (2003), O’Brien et al. (2018), and Fernandes et al. (2019) model interactions that range from mutualistic to parasitic and study host-symbiont coevolution in their interactions with each other, including in the presence of a third species (Gomulkiewicz et al., 2003) and along abiotic gradients (O’Brien et al., 2018). Context-dependent costs and benefits and context-dependent transmission have also been studied in the context of cultural transmission of traits (Ram et al., 2018, 2019).

One of the most important aspects of any symbiosis is how the partners come (or stay) together, i.e., the mode of transmission of the symbiont (Antonovics et al., 2017). For non-context-dependent interactions, the mode of transmission of symbionts is already known to have a strong impact on the distribution and evolution of the symbiosis. In the absence of other feedbacks, symbionts that are transmitted vertically (from parent to offspring) have been found experimentally and theoretically to evolve toward reduce virulence and even mutualism, while symbionts that are transmitted horizontally have been found to evolve intermediate levels of virulence that maximize their transmission (Ewald, 1987; Yamamura, 1993; Ferdy and Godelle, 2005; Stewart et al., 2005; Alizon et al., 2009; Akçay, 2015; Werren et al., 2008; Shapiro and Turner, 2014). Transmission is likely to be important in the evolution of context-dependent symbioses as well, but much less is known about transmission evolution in this context.

Previously, we modeled how the transmission mode evolves in a symbiosis in an environment where the costs and benefits vary in space so that the symbiosis is beneficial to the host in some patches but detrimental in others (Brown and Akçay, 2019). In this setting, the hosts might be expected to evolve a transmission mode that allows them to acquire the symbiont only where it is beneficial and not otherwise. Specifically, one might imagine horizontal transmission to do the trick. However, we showed that achieving such “containment” of the symbiont where it is beneficial non-trivially depends on the eco-evolutionary dynamics of the infection. In particular, containment depends on both the newborn dispersal rate and which component of host fitness the symbiont affects. We showed that populations might evolve fully vertical transmission when the dispersal rate was low and parents’ and offsprings’ environments were highly correlated. Conversely, when the dispersal rate was high, hosts evolved to acquire the symbiont horizontally from neighbors, but only when it affected their lifespan and not when it affected their fecundity. This was because only lifespan effects led to clearing of the infection in patches where it was detrimental so that infection would tend to spread horizontally only where it was beneficial.

In this paper we turn to the question of how temporal in addition to spatial variability in costs and benefits of symbiosis affects transmission mode evolution. Temporally variable environments pose distinct challenges and potential opportunities to hosts. In particular, hosts in a temporally variable environment cannot permanently contain the symbiont to the location where it is beneficial, as this location will change over time. Hosts must now adapt their transmission to allow for fast lost or gain of the symbiont in response to changes in environmental conditions. Another challenge arises in temporally varying environments when environmental conditions are synchronous between locations. In this case, the correlation between parent and offspring environments may be high even with high dispersal rates, potentially increasing the viability of vertical transmission, but also posing a problem by removing a reservoir of hosts with the appropriate infection status for different environmental conditions.

To understand how selection on host transmission strategies may solve these challenges, we model a spatially and temporally variable environment composed of two patches with varying degrees of synchronicity in environmental state. We find that, as in the purely spatial case, hosts can only evolve horizontal transmission when the symbiont affects their lifespan. However, increasing synchronicity allows hosts whose symbiont affects fecundity to rely on vertical transmission of the symbiont even at high newborn dispersal rates. We also find that, unlike the spatial case, pure horizontal or vertical transmission never evolves. Instead, hosts always evolve mixed transmission modes to allow subpopulations to quickly lose and gain the symbiont when the environment changes.

## 2 Methods

### 2.1 The model

We model a population composed of two subpopulations residing in separate patches which can experience different environmental conditions. Each patch contains a fixed number of hosts equal to half the total population. Each time step, a host is chosen to give birth, and, if the newborn survives, an adult in the newborn’s patch is chosen to die. Hosts may be infected with a conditionally mutualistic symbiont whose effect on its host varies depending on the state of the environment. Specifically, the patches experience (potentially different) environments that vary over time between two possible environmental states, State M and State P. In State M, the symbiont benefits its host, increasing its fitness above that of uninfected hosts. In State P, the symbiont is harmful, and infected hosts have lower fitness than uninfected. Each environmental state lasts for a fixed amount of time before changing to the other state. (In the main text we focus on the case where States M and P last for the same amount of time. In the supplement, we consider unequal amounts of time spent in each state, Figures S3 and S4.) We call the time the environment spends in one cycle of State M and State P the time scale of environmental change, sometimes shortened to time scale.

### 2.2 Synchronicity of environmental states

We investigate different degrees of synchronicity between the environmental states in Patches 1 and 2 (Figure 1). At one extreme of the synchronicity continuum, the environmental states in the two patches are completely synchronized, and the two patches are in the same state at all times. On the other end of the spectrum, the patches are completely asynchronous and in opposite states at all times. We refer to the amount by which the state of Patch 1 lags behind the state of Patch 2 as the offset (see Figure 1). For comparison between different time scales of environmental change, we calculate the offset as a fraction of the time scale. An offset of 0 indicates patches are completely synchronous, while an offset of 0.5 indicates complete asynchronicity. An offset of 0.25 indicates that patches are intermediately synchronous and spend half their time in the same state. (We also investigated offsets of 0.125 and 0.375, yielding intermediate results, which we do not show here.)

**Figure 1:**
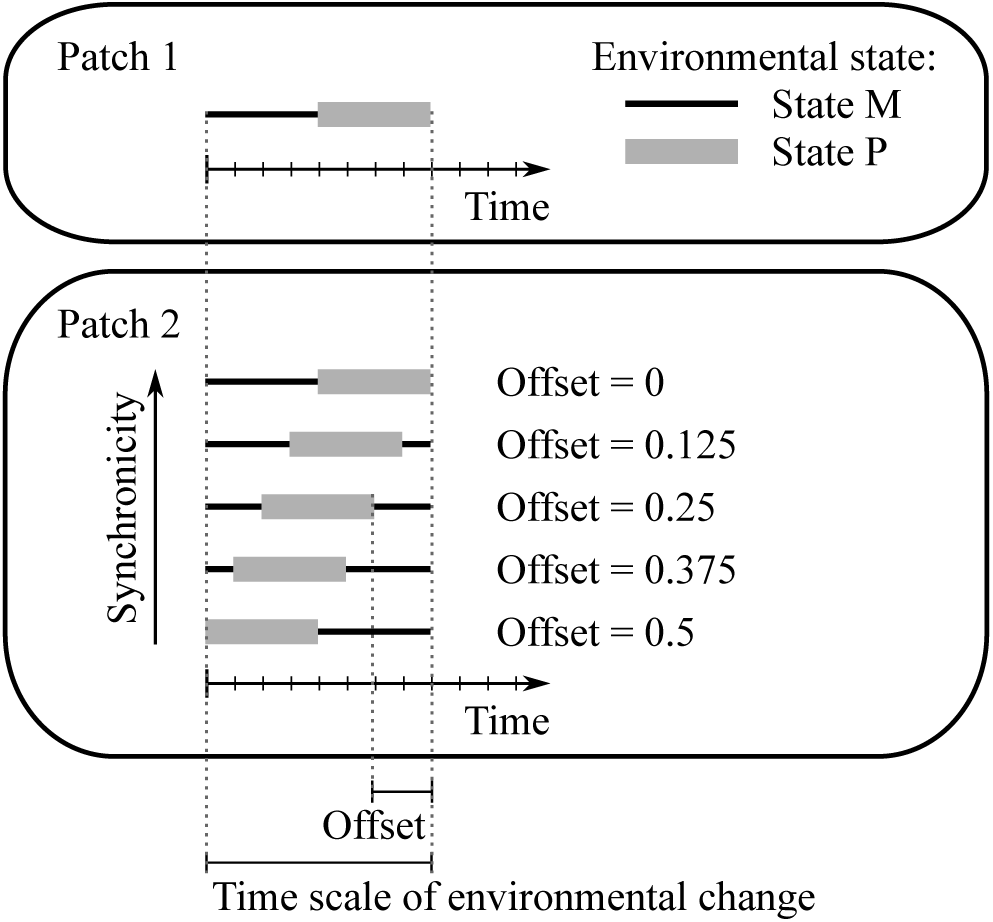
Different levels of synchronicity in environmental state. The offset measures the difference, as a fraction of the time scale of environmental change, between the start of an environmental state in Patch 1 and the start of the same state in Patch 2. An offset of 0 indicates complete synchronicity, while an offset of 0.5 indicates complete asynchronicity. Thin black lines indicate the time a patch spends in State M, and thick gray lines indicate the time it spends in State P.

### 2.3 Symbiont effects on host fitness

We consider two possible ways the symbiont may affect its host. First, the symbiont may affect its host’s lifespan, increasing infected hosts’ lifespan relative to uninfected hosts in State M, and decreasing it in State P. In the main text, we model effects on lifespan through adult host mortality. Hosts with the beneficial infection status for the current environmental state have a lower chance of dying when a newborn host settles in their patch. This is similar to the case where the symbiont affects lifespan through newborn host survival, which we show in the supplement (Figures S1 and S2). The other possibility is that the symbiont affects host fecundity. We model this as increasing the chances that a host with a beneficial infection status for its environmental state will be chosen to reproduce in any given time step.

We model base fecundity, *F*, and mortality, *M*, for each host as variables that depend on their infection status and the environmental state. The chance a particular host gives birth or dies is determined by its base fecundity or mortality compared to the rest of the population (fecundity) or patch (mortality). For a single time step, the probabilities are

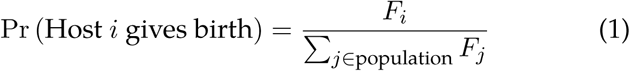

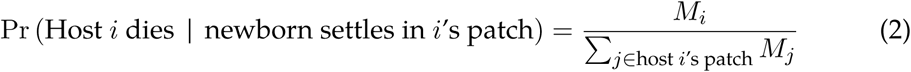

When the symbiont affects host lifespan, hosts with the harmful infection status (hosts uninfected in State in M or infected in State P) have base mortality *M* = 1, while hosts with the beneficial infection status have base mortality *M* = *m* < 1. We set *m* = 0.5. When the symbiont affects fecundity, we set the parameters similarly, except that it is beneficial to have a higher fecundity, so hosts with the beneficial infection status have base fecundity *F* = 1. Hosts with the harmful infection status have fecundity *F* = *f* < 1. We set *f* = 0.5. When the symbiont does not affect an aspect of fitness, we set all base values for that aspect to 1.

### 2.4 Transmission and dispersal

Symbionts in our model can be transmitted between hosts vertically or horizontally. Each host has genetically determined horizontal and vertical transmission probabilities that give their probability of acquiring the symbiont via each mode of transmission. To understand the optimal transmission combination in the absence of biological constraints on transmission, we allow the transmission probabilities to mutate independently and to be any combination of values in the region [0, 1]^2^.

After a newborn is born and has possibly mutated, if its parent is infected, it has a chance to acquire the symbiont via vertical transmission from its parent. After this, the newborn has a chance to disperse to the other patch. We model the dispersal rate as the probability that the newborn leaves its natal patch. At a dispersal rate of 0 the patches behave as separate populations, and at 0.5, the newborn has an equal chance of being in either patch.

If the newborn is still uninfected after it arrives in its final patch, it has a chance to become infected via horizontal transmission. We select a random neighbor in the patch, who, if infected, has a chance to infect the newborn. To prevent the symbiont from being permanently lost from the population, newborns also have a small chance (0.5%) of becoming spontaneously infected if they fail to get infected horizontally. We assume infection can only happen when individuals are newborns and do not allow the newborn to infect adults or adults to infect each other.

### 2.5 Measures of ecological and evolutionary outcomes

We are interested in the transmission mode hosts evolve and how this relates to the ecological dynamics of infection. To understand evolutionary dynamics, we calculate the average horizontal and vertical transmission probabilities after the population has been allowed to evolve for a long period of time (10^6^ generations).

To understand the ecological dynamics, we find the average fraction of infected hosts in each environmental state over a whole number of cycles of environmental change. As hosts are likely to benefit most from transmission probabilities that generate a high fraction of infected hosts in State M and a low fraction in State P, we also calculate the difference between the average fraction of infected hosts in State M and the average fraction infected in State P. We call this difference the symbiont containment.

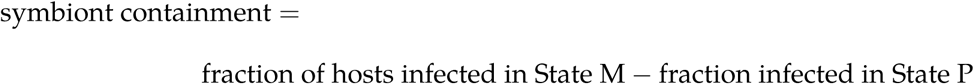

### 2.6 Simulations and numerical analysis

We simulated a context-dependent symbiosis in Julia version 0.6.2 and 0.7.0 (Bezanson et al., 2017). Code for the simulations was adapted from Brown and Akçay (2019) and is available on Github (https://github.com/brownal/ConditionalMutualistsSpaceTime). We simulated a population of 200 hosts, initiated with 50% infection in each patch. To understand transmission evolution, we ran simulations for 2 ⋅ 10^7^ time steps (10^6^ generations). We started simulations at a grid of 9 initial transmission probabilities, with the horizontal and vertical transmission probabilities each either 0, 0.5, or 1. We ran 3 replicate simulations for each set of initial conditions, and calculated the average horizontal and vertical transmission probabilities at the end of each simulation.

We also simulated the ecological dynamics of infection without transmission evolution using a similar procedure. In this case, we ran simulations for 5 cycles of environmental change and found the average infection over the last cycle. We started simulations from a grid of 121 evenly spaced initial transmission probabilities, spaced 0.1 apart. We ran 5 replicate simulations for each set of initial conditions.

We analyzed all simulations using the package pandas (McKinney, 2010, v 2.3.4) in Python version 2.7 (Rossum, 1995) and plotted results in Mathematica v 11 (Wolfram Research Inc., 2017).

We also numerically calculated symbiont infection dynamics for fixed transmission probabilities using Mathematica v 11. Code for this is also available on Github. In this case, we assumed an continuous time and calculated the expected fraction of infected hosts in each state. We did not allow recurrent infection in the numerical analysis, as the infection cannot be lost by chance in this case.

## 3 Results

### 3.1 Symbiont containment

#### Symbiont affects host lifespan

We simulated infection dynamics at fixed transmission probabilities to determine the effects of transmission on containment. When the symbiont affects host lifespan and patches experience environments that are completely synchronized, containment is highest in a band of transmission probabilities where horizontal and vertical transmission probabilities trade off against each other (Figure 2). At intermediate transmission probabilities, the band curves up somewhat, toward slightly higher total transmission giving the highest containment. Interestingly, the band of high containment does not include 100% horizontal or vertical transmission, both of which lead to very low containment. At lower levels of synchronicity, the pattern is similar when the dispersal rate is low. As the dispersal rate increases, containment decreases at high vertical transmission probabilities. Containment is then highest when vertical transmission is low, but not absent, and the horizontal transmission probability is high, but still less than 1. These results are counter to the case with purely spatial heterogeneity, where complete vertical or horizontal transmission by itself can generate containment.

**Figure 2:**
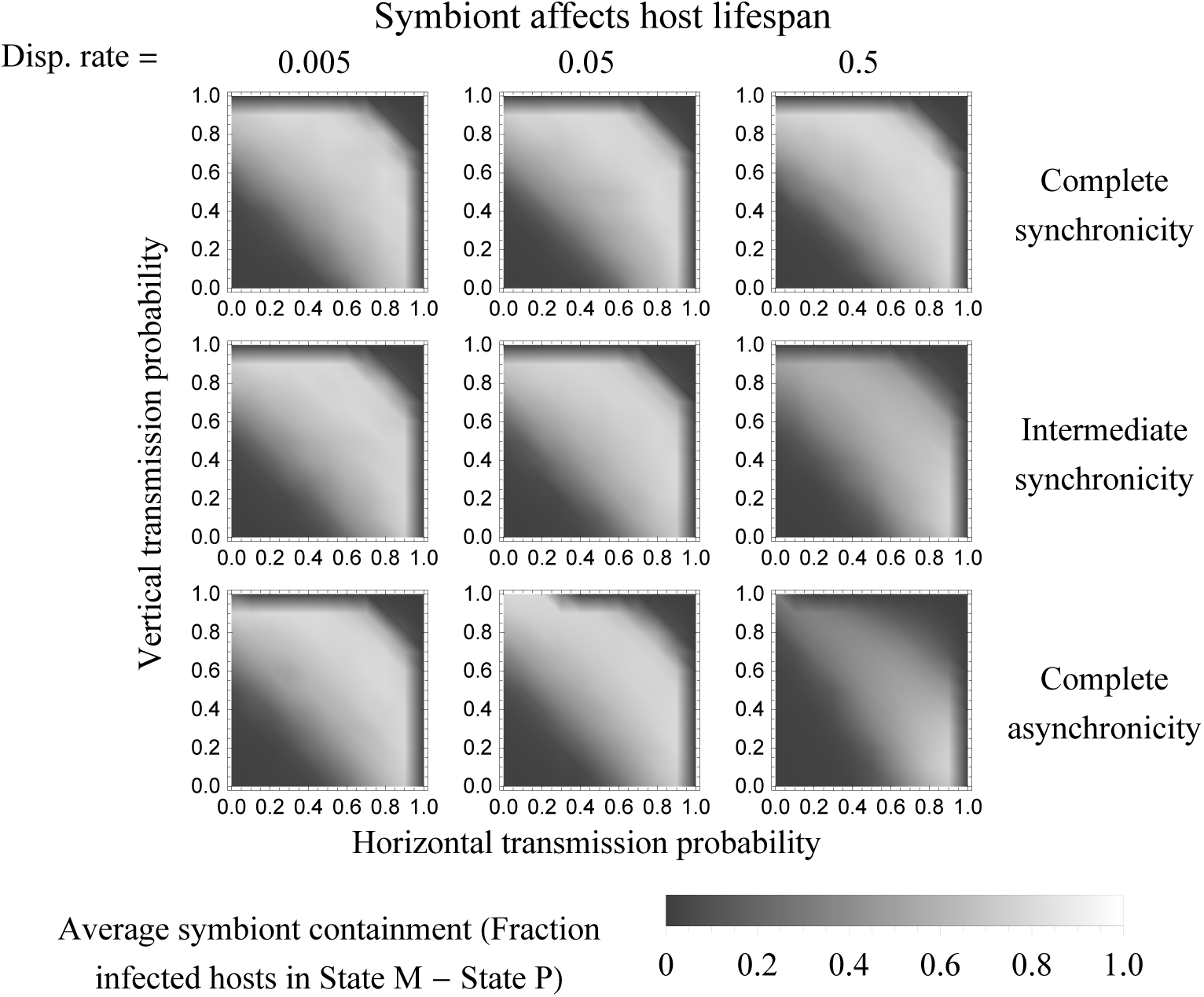
Average symbiont containment with fixed transmission probabilities when the symbiont affects host lifespan. Simulations were run for a grid of fixed transmission probabilities spaced 0.1 apart. Time scale of environmental change: 160 generations (32,000 time steps). Simulations run for 6 cycles of environmental change (192,000 time steps), with the fraction of infected hosts at 1000 evenly spaced time points in the last cycle (32000 time steps) used to find the average infection in each environmental state. Containment was determined by subtracting these average infection levels. Containment was then averaged across 5 replicate simulations.

#### Symbiont affects host fecundity

When the symbiont affects fecundity, containment decreases at high horizontal transmission probabilities (Figure 3). Containment is generally highest at high vertical transmission probabilities and low or moderately low horizontal transmission probabilities. However, 100% vertical transmission generally leads to low containment, as does 0% horizontal transmission. When the patches are synchronized, the maximum containment is similar to when the symbiont affects lifespan. On the other hand, when patches are asynchronous, containment decreases for all transmission probabilities with increasing dispersal. At complete asynchronicity and a dispersal rate of 0.5 (parent and offspring environments are completely uncorrelated), containment is essentially 0 for all transmission probabilities.

**Figure 3:**
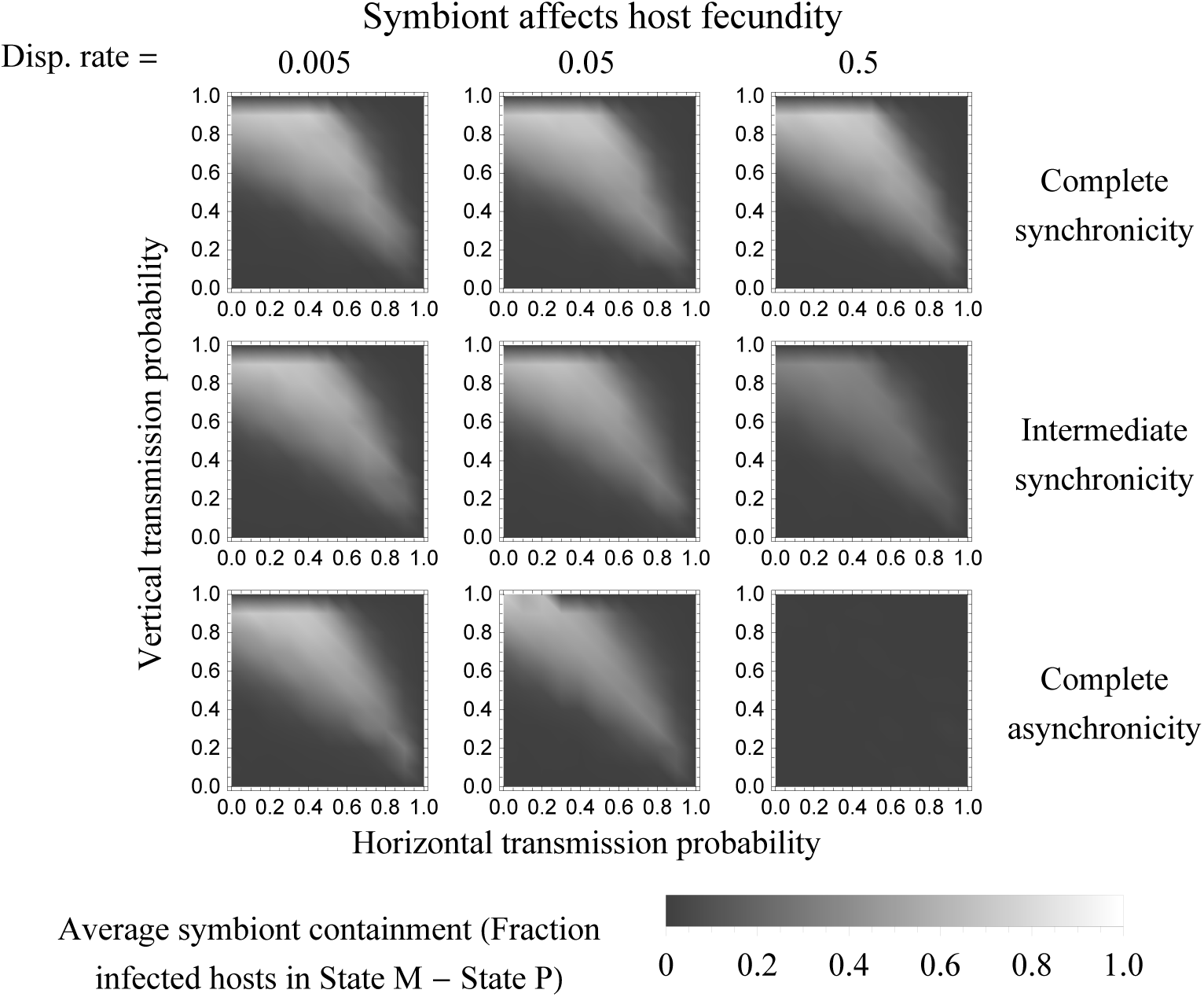
Average symbiont containment with fixed transmission probabilities when symbiont affects host fecundity. All other parameters the same as Figure 2. In some cases where containment was generally low, spontaneous infection occasionally produced negative containment; shown here as zero containment (see SI for further discussion).

#### Analytical model

Our analytical model of containment, evaluated numerically, confirms these results. One exception is that because the analytical model allows for the loss of the symbiont in a finite population, some containment is possible when the horizontal or vertical transmission probability is 1 (Figure S7). Also, due to the lack of spontaneous infection, the infection can be lost in some cases where it remained in the simulations. This largely affects the case where the two patches experience synchronized environments and the symbiont affects host lifespan. In this case, while the region where infection can be maintained is much smaller, a curve representing a trade-off between horizontal and vertical transmission still produces the highest containment.

### 3.2 Transmission mode evolution

The vertical and horizontal transmission rates evolve toward values that result in high containment (Figure 4).

**Figure 4:**
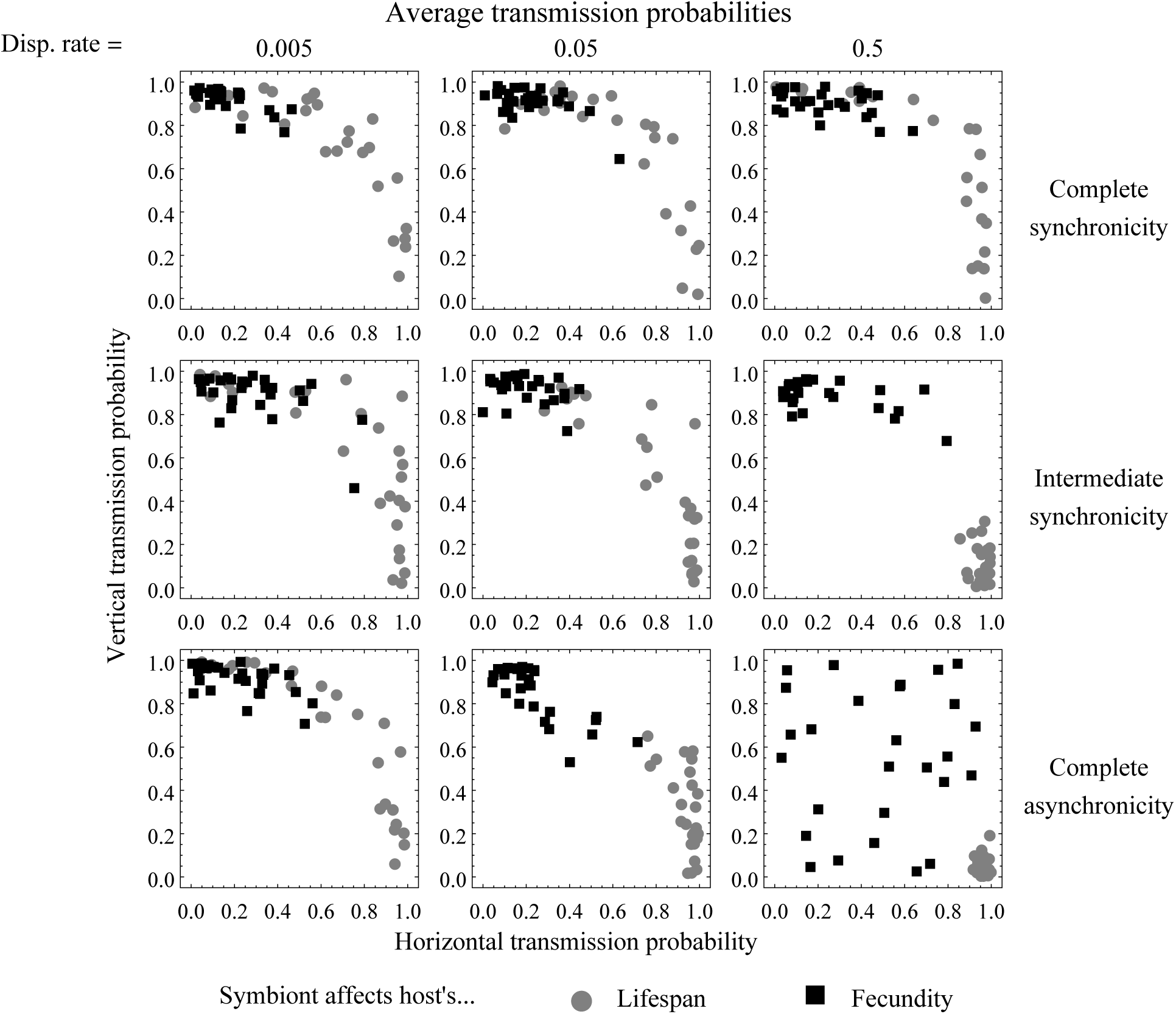
Average transmission probabilities after evolution. Each point shows the average horizontal and vertical transmission probabilities for a single simulation after 10^5^ generations (2 ⋅ 10^7^ time steps) of transmission evolution. Time scale of environmental change = 160 generations (32,000 time steps). Simulations were started from a grid of transmission probabilities spaced 0.5 apart. 3 replicate simulations were run from each starting point.

#### Symbiont affects host lifespan

When the symbiont affects host lifespan and both patches are synchronized in time, transmission mode evolves to be somewhere in the band of approximately equal horizontal and vertical transmission that leads to high containment. This creates the appearance of a trade-off between the two transmission modes, even though both transmission probabilities could evolve completely unconstrained by each other. Within this band of high containment, the transmission probabilities appear to evolve neutrally. Asynchronicity does not break this apparent neutrality under low dispersal, when the dynamics resembles the synchronous case. However, with high dispersal rates and asynchronous environments, hosts evolve high horizontal and low vertical transmission.

#### Symbiont affects host fecundity

When the symbiont affects host fecundity and patches are synchronized, hosts evolve high vertical transmission and low levels of horizontal transmission. The same is true when patches are asynchronous and the dispersal rate is low. As the dispersal rate increases in asynchronous environments and neither vertical nor horizontal transmission can contain the symbiont, more horizontal transmission appears possible, likely because transmission appears to be evolving neutrally.

### 3.3 Effect of time scale of environmental change

The time scale over which environmental change occurs also affects both symbiont containment and transmission evolution. In particular, the results highlighted above only occur at intermediate time scales. At very short of time scales of 800 (each patch remains in its state for 2 generations) and 80 (each patch remains in its state for 0.2 generations), transmission probabilities evolve neutrally (Figure S5), while at very long time scales of 8 ⋅ 10^6^ time steps, transmission evolution resembles the case where there is purely spatial variation in symbiont quality (Figure S6).

## 4 Discussion

Conditionally beneficial symbionts in temporally varying environments present a challenge for their hosts, which would benefit from acquiring the symbiont only when and where it is beneficial. Our results show that hosts have a range of options for achieving such containment of symbionts. Generally, the evolutionary outcomes for each transmission mode depend on the aspect of fitness the symbiont affects (for horizontal transmission) and the synchronicity of environmental states across space (for vertical transmission). When the symbiont conditionally affects host lifespan but not fecundity, hosts are able to evolve horizontal transmission and contain symbionts to environments where they were beneficial. When the symbiont affects host fecundity, hosts with the “wrong” infection status tend to remain in the population, as they are no more likely to die than hosts with the beneficial infection status. Under this scenario, horizontal transmission causes newborns to acquire this harmful infection status from their neighbors, and thus is selected against.

In contrast, we find that vertical transmission can always evolve at low newborn dispersal rates. At high dispersal rates, correlation between parent and offspring location is reduced, and vertical transmission does not evolve in asynchronous environments. Synchronicity in environmental state across space “rescues” vertical transmission by creating a correlation in environmental conditions regardless of location. When the symbiont affects host lifespan, in many cases, hosts can use either horizontal or vertical transmission to achieve symbiont containment. In these cases, containment can also be achieved with a range of mixtures of the two transmission modes.

Strikingly, the range of mixtures of horizontal and vertical transmission embodies an apparent trade-off between the two transmission modes even though we do not impose any direct costs or enforce any constraints. This trade-off arises in our model purely due to the eco-evolutionary feedback with infection dynamics that results from each combination of transmission probabilities. Specifically, increasing both transmission modes together beyond a certain level precludes hosts from losing the symbiont when it becomes harmful. Thus, adaptive evolution tends to lower one transmission probability if the other one increases. Within this “adaptive band” of mixed mode transmission probabilities, fitness is nearly neutral. This emergent tradeoff due to eco-evolutionary feedbacks has potential implications for local adaptation and differentiation of populations. It suggests that populations that independently evolve under similar environmental conditions and with the exact same symbiont may still experience reproductive isolation if they converge on different points in the band of adaptive transmission probabilities. In that case, hybrid lineages between the populations could end up with transmission probabilities outside the adaptive band and thus with lower fitness than their parent lineages, leading to reproductive isolation.

Another potential outcome for hosts in the long term is that physiological or developmental mechanisms that canalize the trade-off between horizontal and vertical transmission might be selected for to increase hosts’ chances of remaining along the trade-off line in the face of mutations in transmission (Rice, 2008; Watson et al., 2014; Pavličev and Cheverud, 2015). Many symbioses do exhibit some physiological constraints on transmission, and it would be interesting if, in some cases, this were a consequence of evolutionary trade-offs, rather than a cause of them.

Our results stand in interesting contrast to the pure spatial heterogeneity case we analyzed before (Brown and Akçay, 2019). With temporal heterogeneity, we find that purely vertical or purely horizontal transmission mode does not lead to high containment, whereas in the pure spatial heterogeneity case only pure horizontal or pure vertical transmission could achieve maximum containment (Brown and Akçay, 2019). This is in part because under temporal heterogeneity, hosts are periodically selected to quickly lose the symbiont, which cannot happen if either transmission probability is very close to one. Conversely, having both types of transmission but at less than maximum probability, on the other hand, allows the symbiont to spread between lineages (horizontal transmission) and locations (vertical transmission) when it becomes beneficial. Temporal variability therefore increases the benefits of mixed-mode transmission for hosts and even produce an emergent trade-off between horizontal and vertical transmission.

Our results apply to the case where temporal variation happens on intermediate time scales. If the environment changes much more quickly than the ecological dynamics of infection, neither parents’ nor neighbors’ infection status can predict whether the symbiont is beneficial. In this case, transmission evolves neutrally. When the environment changes more slowly than host evolution, hosts can evolve as if the symbiont’s effects purely varied in space. Our results only apply when the environment changes more slowly than ecological dynamics but more quickly than evolutionary ones.

There are several examples of symbioses that likely fall within this regime. The fungus *Epichloë amarillans* increases the fecundity of its hosts, the grass *Agrostis hyemalis* under dry conditions, while decreasing host biomass, which may affect lifespan or future fecundity, in the presence of soil microbes (Davitt et al., 2011). The fungus is both vertically and horizontally transmitted in this host, though transmission in the studied population is predominantly vertical (Davitt et al., 2011). As the authors find no environmental effects on transmission, they suggest that the differences in endophyte prevalence in natural populations may be due to the environmental context. As water availability likely varies asynchronously (as both wet and dry locations exist simultaneously), our model would suggest symbiont containment if host dispersal is low. The fact that high, but not complete, vertical transmission evolves in response to effects on fecundity also matches with the predictions of our model.

The aphid *Aphis craccivora* also experiences a conditional mutualism with the bacterial symbiont *Arsenophonus*. *Arsenophonus* improves aphid ability to eat locust at the expense of other plants, in particular increasing aphid population growth on locust while decreasing it on alfalfa, suggesting that the bacterium at least conditionally affects fecundity (Wagner et al., 2015). The symbiont is also vertically transmitted, and horizontal transmission may also be possible, based on evidence from related hosts and similar symbiont species (Russell and Moran, 2005; Wagner et al., 2015). The exact location of locust trees probably changes over timescales intermediate between ecological and evolutionary dynamics. There is evidence of symbiont containment to aphid populations found on locust plants (Brady and White, 2013). With vertical transmission and fecundity effects, we should expect symbiont containment if dispersal is low or environmental synchronicity is high. While the exact spatial distribution of locust trees probably varies in different environments, parent-offspring correlation may be high if aphids selectively disperse to host plants of the same species as their natal plant. As some aphids were specialized on their alfalfa food source in the absence of infection, there is some evidence for prolonged association of a lineage with a specific type of host plant.

Another context-dependent interaction with aphids is that of pea aphids, *Acyrthosiphon pisum*, with their bacterial symbiont *Hamiltonella defensa*, which protects them from parasitoid wasps but increases their predation by ladybugs (Polin et al., 2014). Since the presence of predators and parasitoids should vary in time, this makes *H. defensa* a conditional mutualist whose effects vary over time. If this is the case, we would expect to observe imperfect transmission. Because this symbiosis has a lifespan cost, at least some horizontal transmission would seem likely, with horizontal transmission becoming more likely as the correlation between the predators/parasitoids around parents and offspring decreases. Interestingly, vertical transmission of *H. defensa* has been found in this symbiosis, with extremely consistent vertical transmission when *A. pisum* reproduces clonally (Moran and Dunbar, 2006). However, there is the potential for vertical transmission failures during sexual reproduction (Peccoud et al., 2014) or post-coinfection with another secondary symbiont (Rock et al., 2018). Horizontal transmission of *H. defensa* in *A. pisum* has not yet been observed, but evidence of horizontal transmission of *H. defensa* in other aphid species exists (Gehrer and Vorburger, 2012; Li et al., 2018). Kwiatkowski and Vorburger (2012) modeled the ecological dynamics of this symbiosis in a spatially homogeneous environment and found that high (but not complete) vertical transmission combined with low horizontal transmission can maintain the symbiont at intermediate frequency in the population (as does *H. defensa*-infected hosts paying an induced cost when attacked by a parasitoid). Our results suggest a mechanism by which hosts might evolve to maintain this type of transmission, which also leads to symbionts at intermediate frequencies in a spatially heterogeneous population.

Our model makes predictions for the outcome of context-dependent symbioses in environments that vary in both space and time, but there are still questions that remain. In particular, it is not clear how symbionts will evolve in a context-dependent symbiosis when the environment varies over time. Understanding symbiont evolution in such symbioses and in response to selection for transmission trade-offs in hosts may give more insight into the host-symbiont coevolution and the eventual fate of these symbioses. Furthermore, some hosts, such as aphids, have been found to engage in multiple context-dependent interactions, and many other context-dependent interactions have effects that changed based on the presence of a third symbiont. Understanding how multiple symbionts affect host evolution, and how host response to infection feeds back on symbiont dynamics, will help us understand the effects of context-dependence on larger ecological dynamics.

Our results also suggest that understanding the temporal synchronicity of environmental conditions across space is needed to predict transmission evolution. Measuring this in natural populations of hosts in context-dependent interactions could allow for predictions of transmission evolution in these symbioses. Along with that, as natural environments likely change stochastically over time, extending the model to different types of environmental change over time will be helpful for making accurate predictions for existing symbioses.

Our model predicts that temporal environmental variation can strongly affect transmission evolution in context-dependent symbioses. In particular, synchronicity in environmental conditions across space can rescue vertical transmission as a method of symbiont containment, when symbiont effects on fecundity make horizontal transmission impossible. Our model also suggests an emergent trade-off between horizontal and vertical transmission, suggesting that physiological constraints on transmission are not necessary for a trade-off between transmission modes to evolve. Furthermore, our model also suggests that mixed-mode and imperfect transmission could benefit hosts, by allowing them to respond to environmental changes.

## Acknowledgments

AB was funded by a DOD NDSEG fellowship. Computational work was supported in part by a NAS Keck Futures Initiative grant to J Van Cleve, EA, and T Linksvayer.

