## Supplementary figures for "Evolution of symbiont transmission in conditional mutualisms in spatially and temporally variable environments"

May 5, 2020

#### 1 Symbiont affects lifespan via newborn establishment

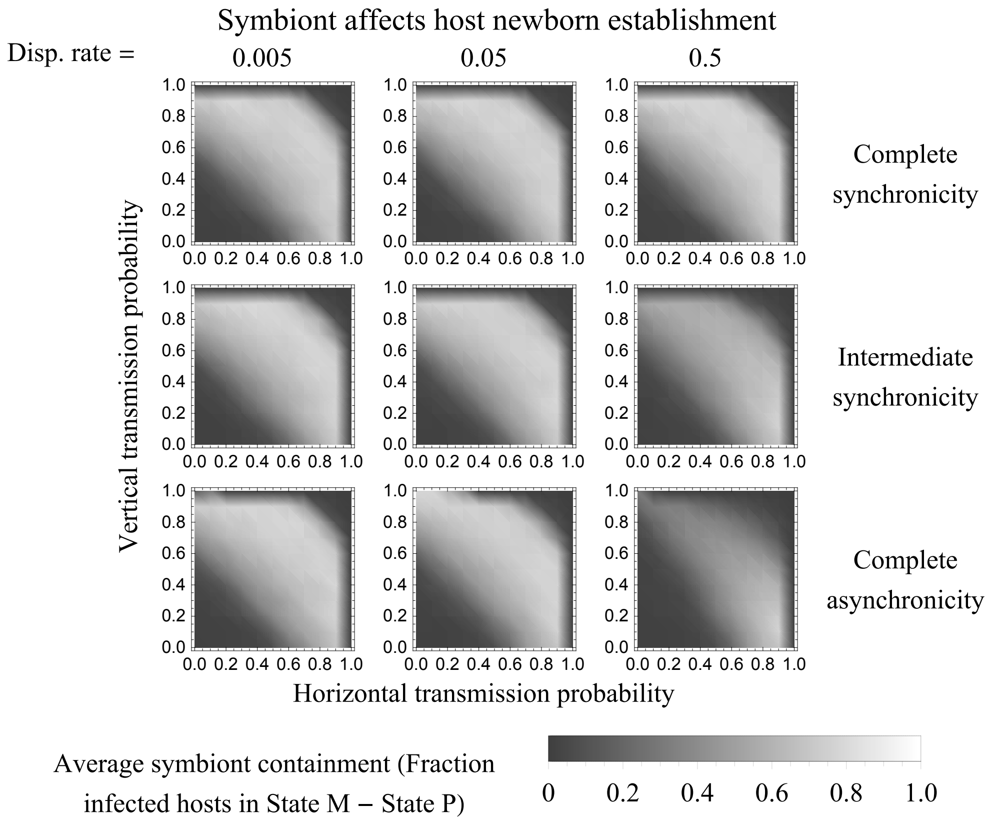


Figure 1: Average symbiont containment when the symbiont affects host lifespan via newborn host establishment. Simulations were run for a grid of fixed transmission probabilities spaced 0.1 apart. (No transmission evolution, $\mu = 0$). Time scale of environmental change: 160 generations (32,000 time steps). Simulations run for 6 cycles of environmental change (192,000 time steps), with the fraction of infected hosts at 1000 evenly spaced time points in the last cycle (32000 time steps) used to find the average infection in each environmental state. Containment was determined by subtracting these average infection levels. Containment was then averaged across 5 replicate simulations. Containment is similar to that shown in the main text when the symbiont affects lifespan via adult host mortality.

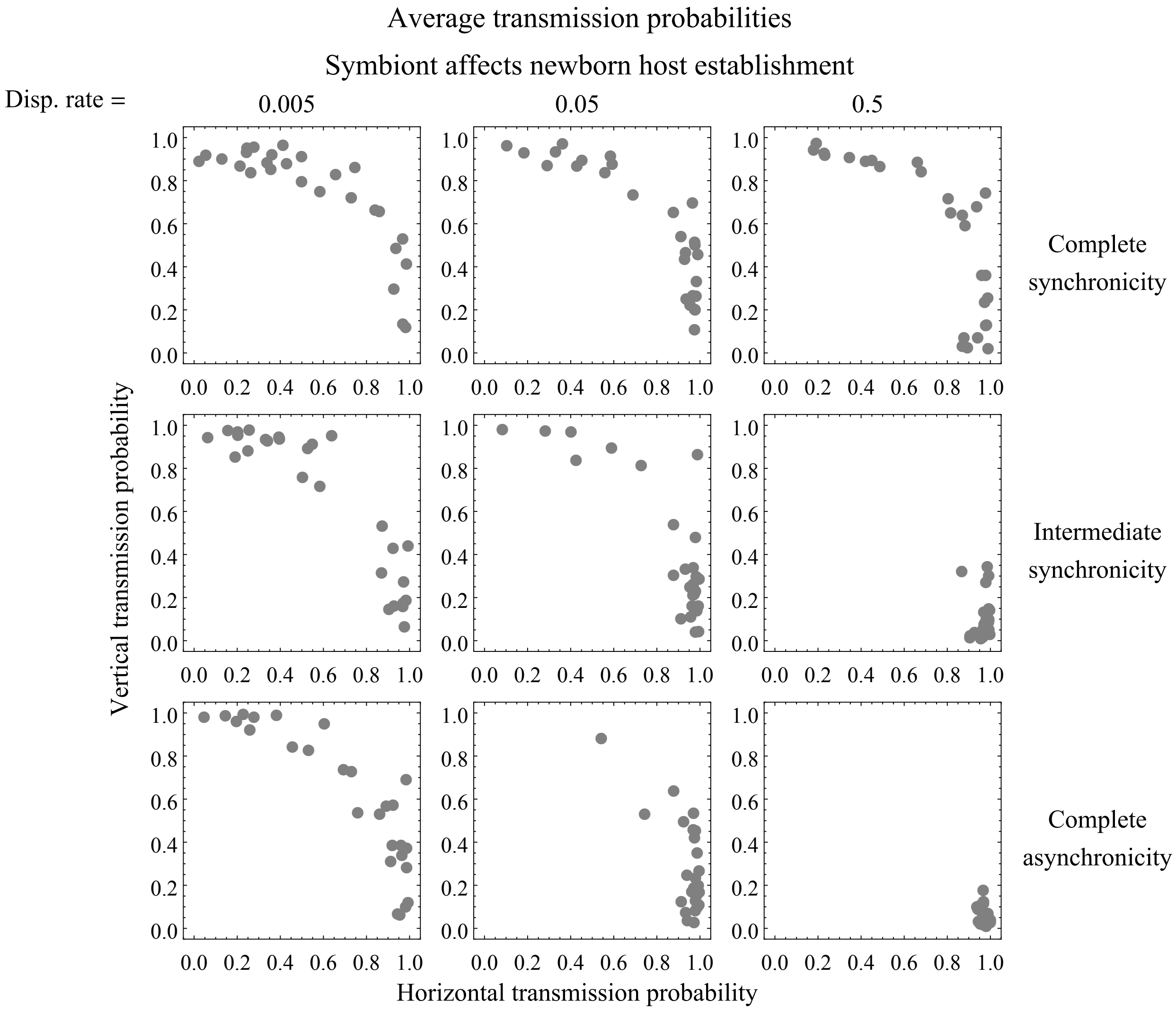


Figure 2: Average transmission probabilities after evolution when the symbiont affects lifespan via via newborn host establishment. Each point shows the average horizontal and vertical transmission probabilities for a single simulation after $10^5$ generations ($2 \cdot 10^7$ time steps) of transmission evolution. Time scale of environmental change = 160 generations (32,000 time steps). Simulations were started from a grid of transmission probabilities spaced 0.5 apart. 3 replicate simulations were run from each starting point. Average transmission is similar to that shown in the main text when the symbiont affects lifespan via adult host mortality.

### 2 Patches spend unequal amounts of time in M and P states

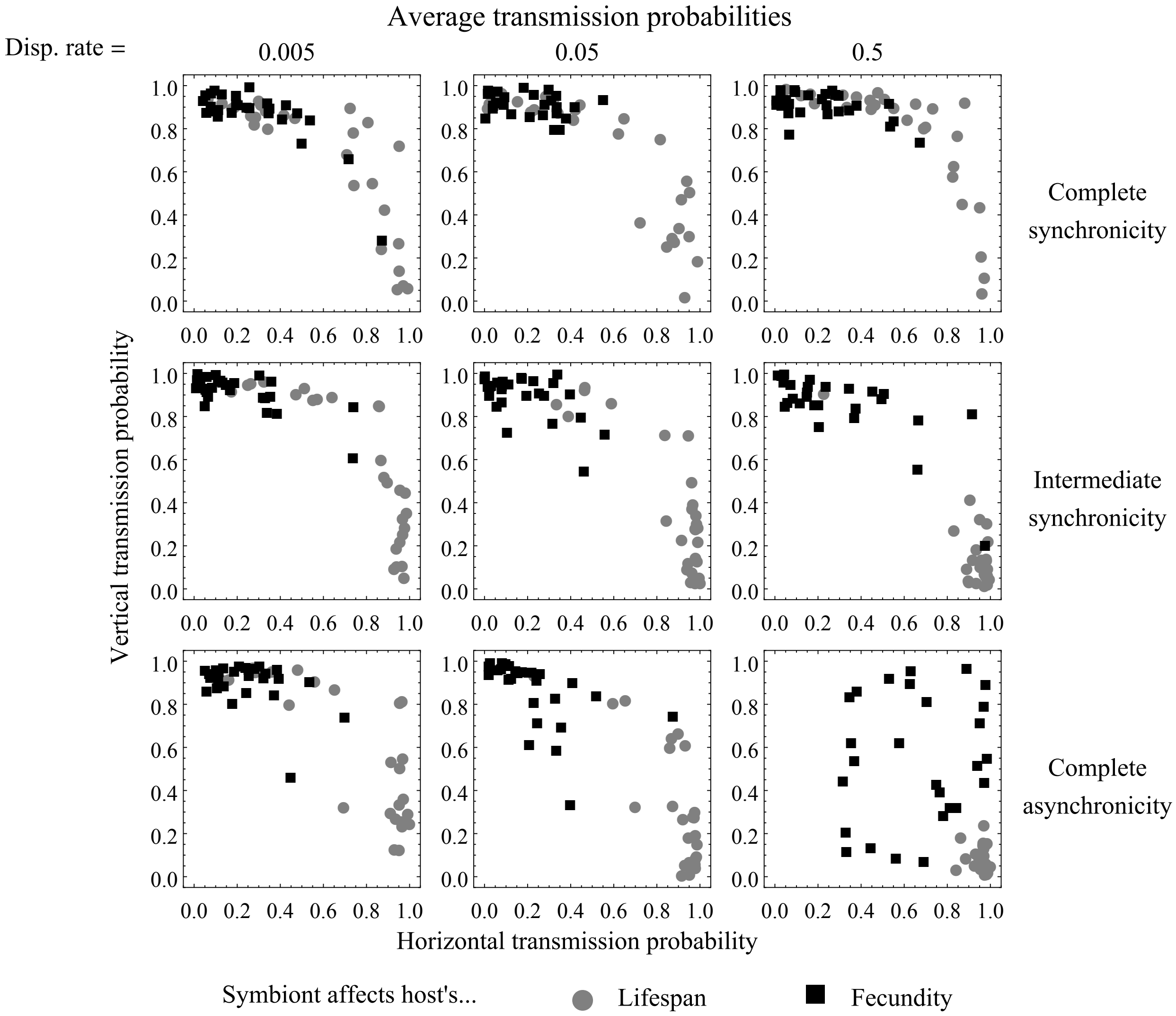


Figure 3: State M is more common than State P. Transmission evolution is similar to the case where patches spend the same amount of time in each state. Plots show average transmission probabilities after 80,000 generations of evolution ($1.6 \cdot 10^7$ time steps). Patches are in State M 62.5% of the time and State P 37.5% of the time. Other parameters the same as Main Text Figure 4.

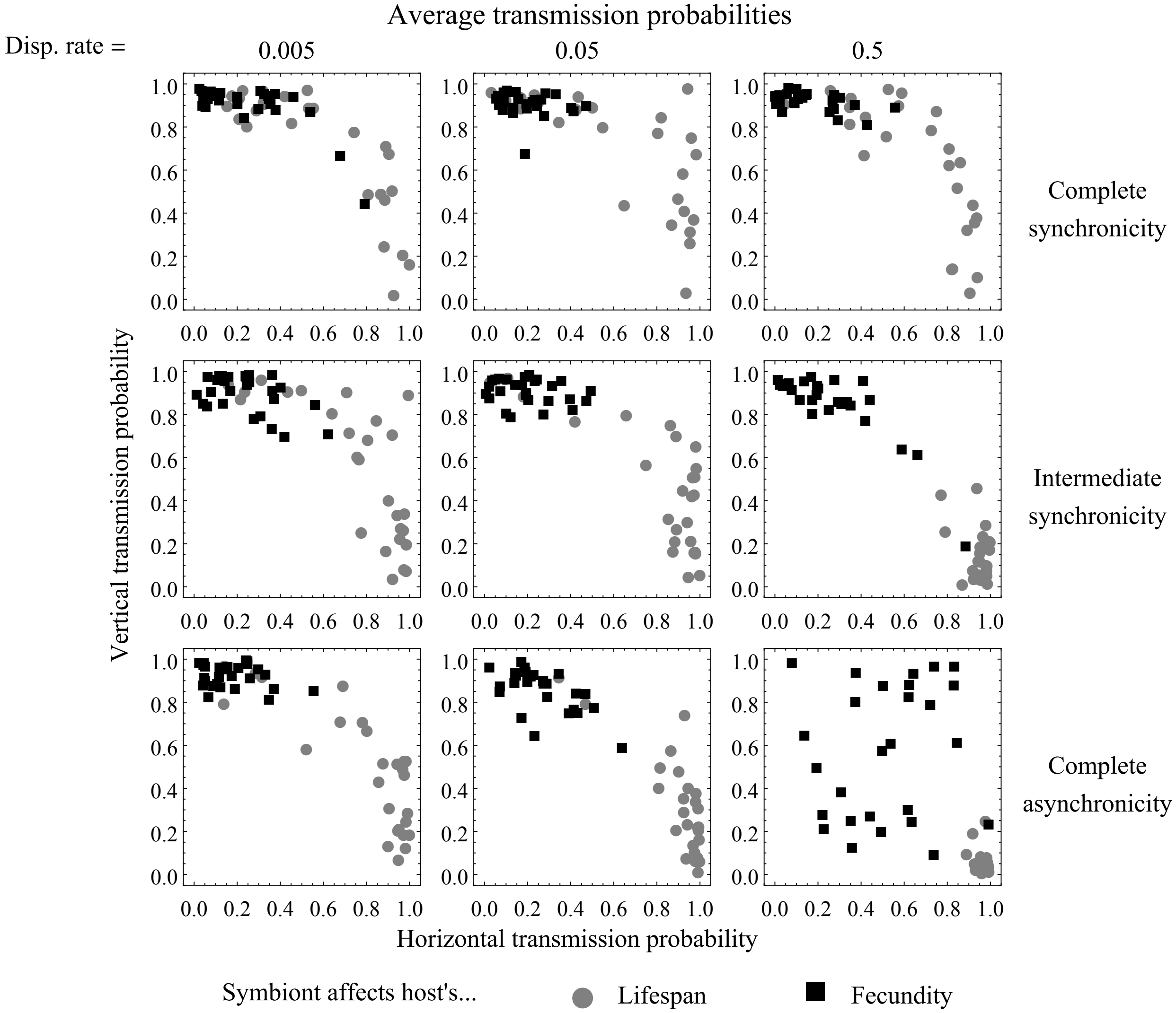


Figure 4: State P is more common than State M. Transmission evolution is similar to the case where patches spend the same amount of time in each state. Plots show average transmission probabilities after 80,000 generations of evolution ($1.6 \cdot 10^7$ time steps). Patches are in State M 37.5% of the time and State P 62.5% of the time. Other parameters the same as in Main Text Figure 4.

### 3 Time scale of environmental change is very short

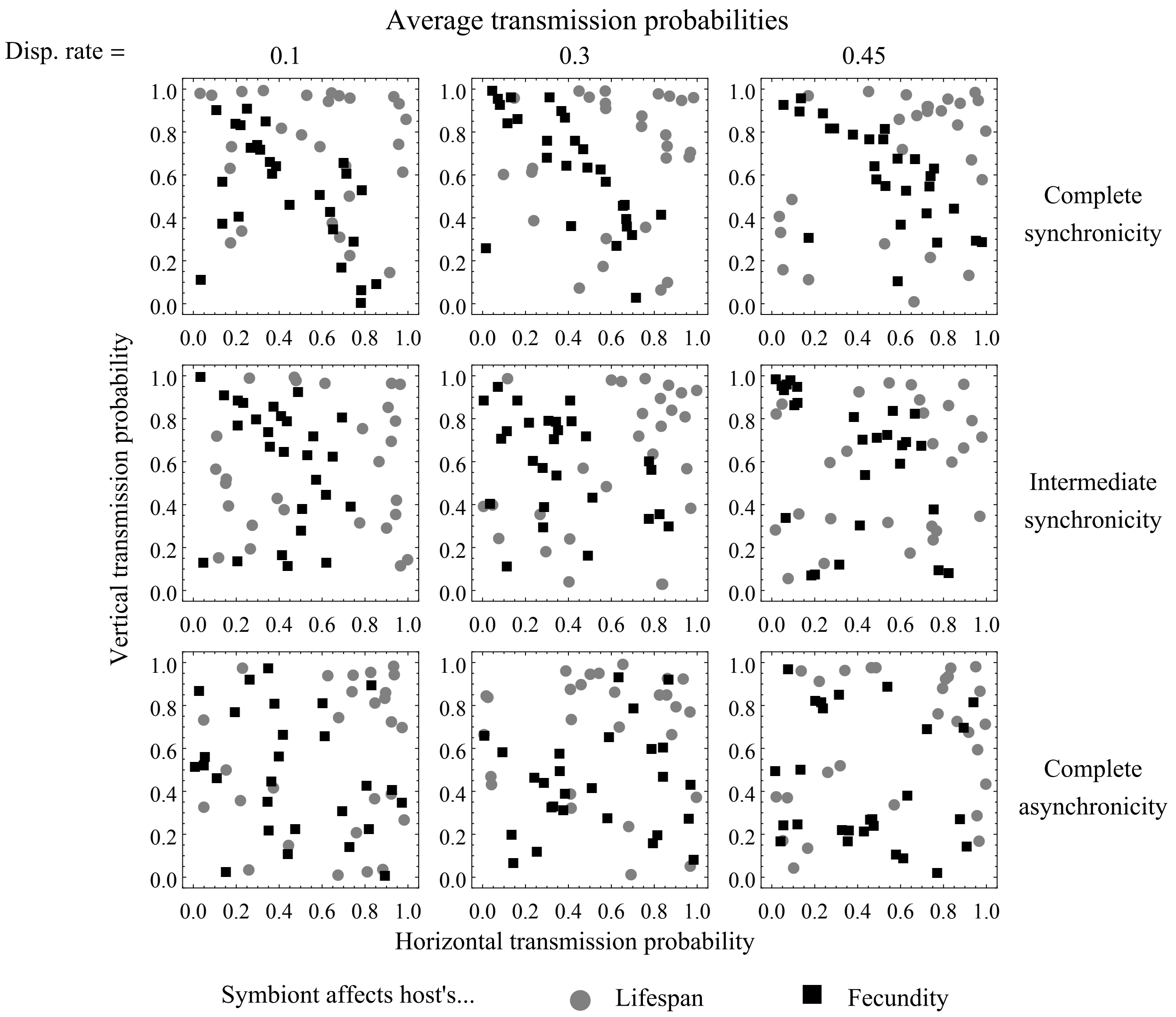


Figure 5: Time scale of environmental change is very short (4 generations, 800 time steps). Transmission evolution appears largely neutral. This is probably because the environment changes so quickly the costs and benefits of infection average out over short time scales, making infection status (and thus transmission) behave like a neutral trait. Plots show average transmission probabilities after 8000 generations of evolution ($1.6 \cdot 10^6$ time steps). Other parameters the same as in Main Text Figure 4.

### 4 Time scale of environmental change is very long

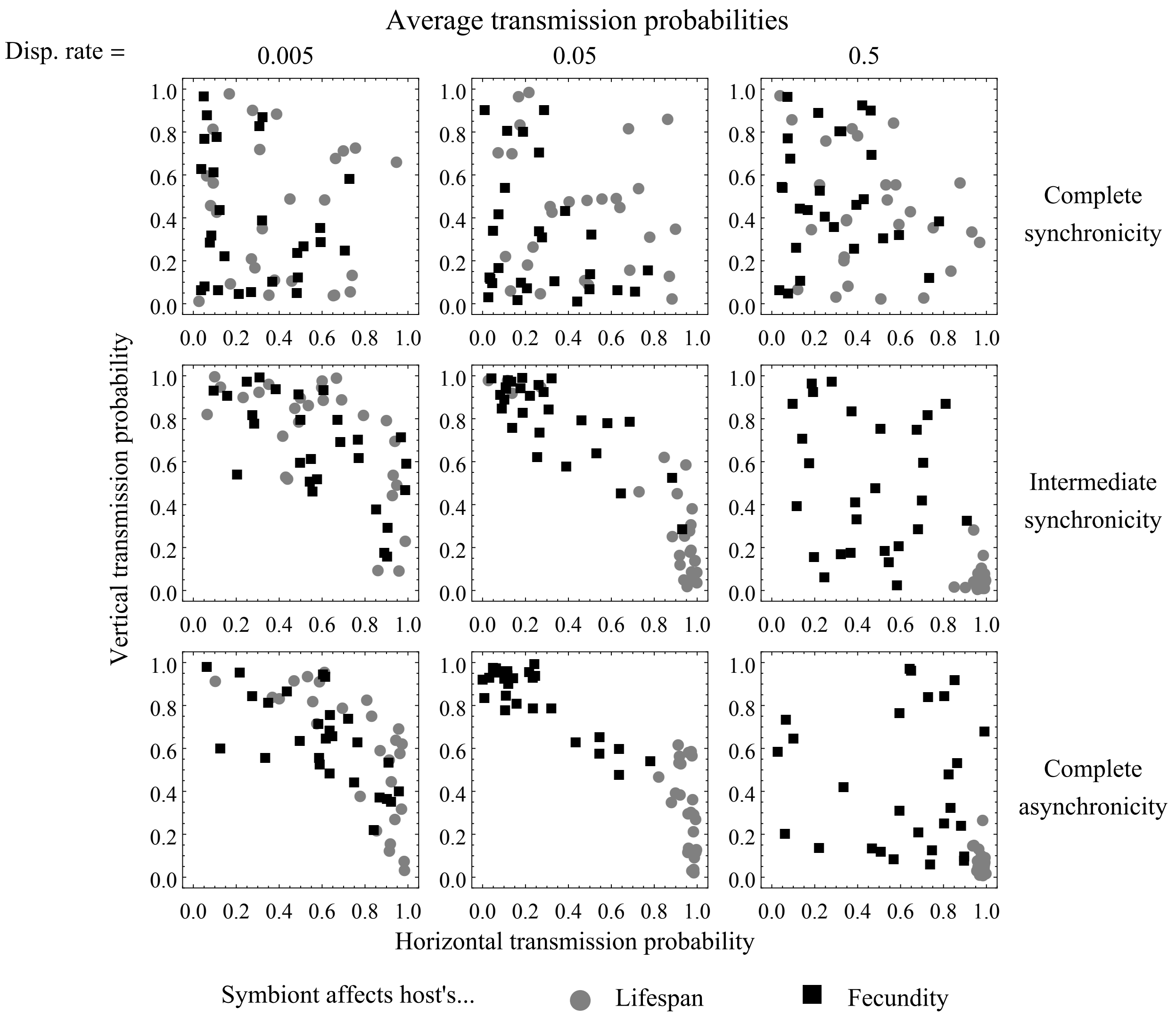


Figure 6: Time scale of environmental change is very long (40000 generations, $8 \cdot 10^6$ time steps). Plots show average transmission probabilities after 80000 generations of evolution ($1.6 \cdot 10^7$ time steps). Other parameters the same as in Main Text Figure 4.

### 5 Results of analytical model of ecological dynamics

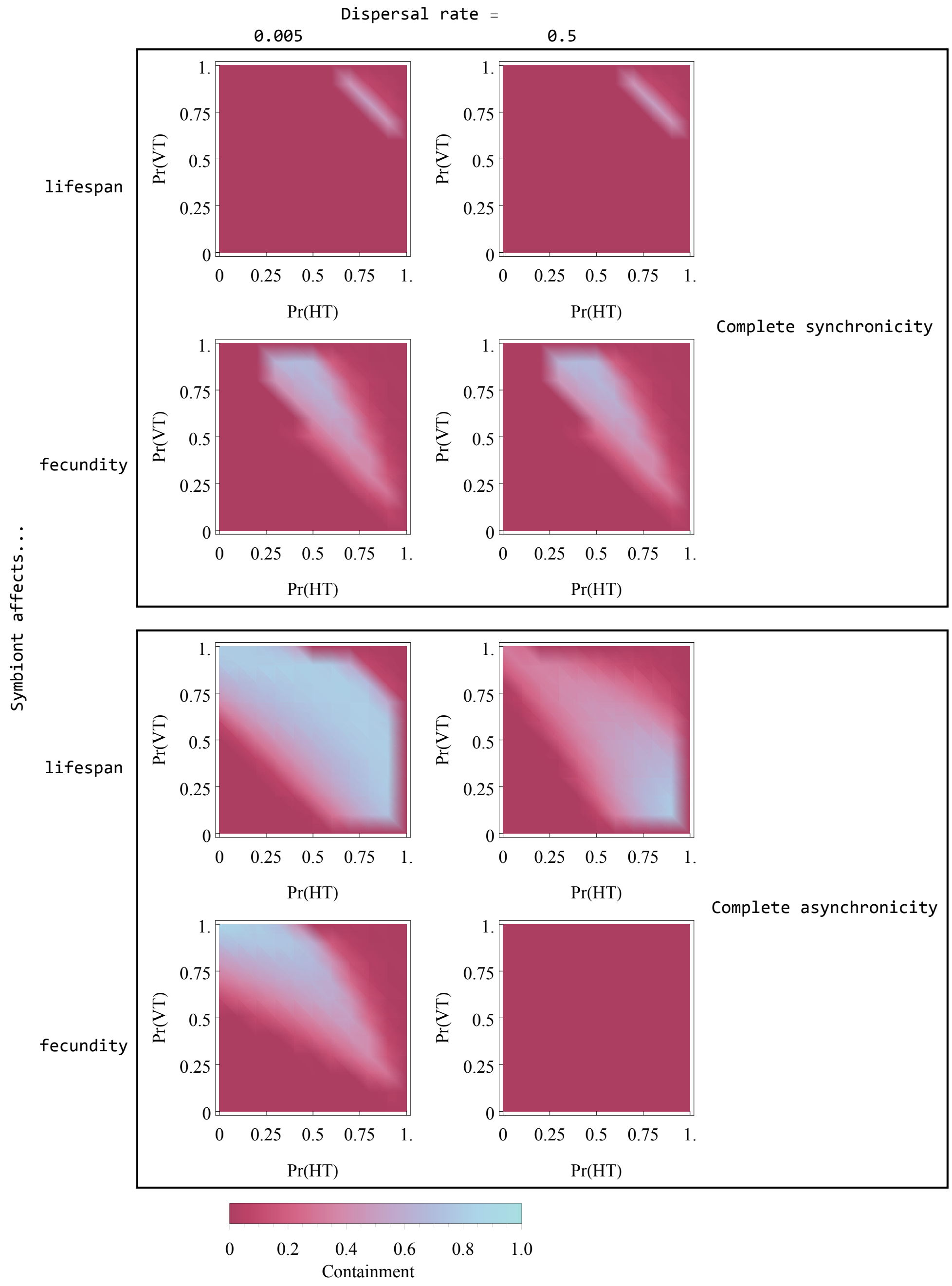


Figure 7: Average containment for analytical model. Infinite population; time scale of environmental change is 160 generations. Average containment was calculated numerically for a grid of horizontal and vertical transmission probabilities spaced 0.1 apart by forecasting the fraction of infected hosts in each patch for 5 cycles of environmental change. The average fraction of infected hosts in each environmental state for the last cycle was used to calculate the average containment. (In one case a longer number of cycles was used; see code.)
